# A Novel, Improved, Application for the Normalization of RNA-seq Expression Data in Complex Polyploids

**DOI:** 10.1101/2021.07.27.453947

**Authors:** Dyfed Lloyd Evans

## Abstract

Much of the work on the normalization of RNA-seq data has been performed on human, notably cancer tissue. Little work has been done in plants, particularly polyploids and those species with incomplete or no genomes. We present a novel implementation of GeTMM (Gene Length Corrected TMM) that accounts for GC bias and works at the transcript level. The algorithm also employs transcript length as a factor, allowing for incomplete transcripts and alternate transcripts. This significantly improves overall normalization. The GCGeTMM methodology also allows for simultaneous determination of differentially expressed transcripts (and by extension genes) and stably expressed genes to act as references for qRT-PCR and microarray analyses.

## Introduction

As RNA-seq analysis has come to dominate transcriptomic analyses (a switch from the previous front-runner of microarrays) due to its massively sequencing of cDNA (Mortazavi et al. 2008). RNA-seq also allows for the study of novel transcripts along with a better range of detection and lower technical variability (Zhao et al 2014) and provides a high degree of agreement with the currently accepted ‘gold standard’ in transcriptomic expression, qRT-PCR at both the absolute and relative expression analysis levels (Su and Mason 2014). A typical RNA-seq experiment involves several steps to enable data analysis: trimming (Williams et a1 2016), alignment (mapping) (Borozan et al. 2013; Yang et al. 2015), read counting, data normalization and analysis (Lin et al. 2015; Li et al. 2017).

As much of the developmental work in RNA-seq analyses derives from the human genome, sequence reads are aligned to a reference genome, and the number of reads mapping to that feature are proportional to the length and abundance of the feature — with the ‘gene’ feature being a surrogate for all the transcripts transcribed from that gene.

Little work has been done on error correction of RNA-seq data for the discovery of stably-expressed transcripts/genes and differentially expressed genes in plants, particularly complex polyploids though there are some whole genome analyses (Park et al 2019; Gupta et al 2012). However, most of the developmental work in this area has been performed on human data, particularly comparing cancer and normal tissue datasets. Typical normalization methods for RNA-seq data typically allow for either intersample comparisons (for differentially expressed genes) or intrasample comparison (for discovery and/or validation of gene signatures) (Smid et al. 2018).

For other organisms, where no genome is available a transcriptome assembly may be substituted for the complete genome. Where both the genome and transcriptome are incomplete reads may be mapped g to a subset that only includes the transcripts/genes of interest (Peri et al 2020). However, as depth of RNA-seq data can vary, normalization has to be performed to correct for differences between sequencing runs (e.g. library size and relative abundances) prior to any downstream analyses.

The most commonly used RNA-seq normalization methods are TMM, as implemented in edgeR (Robinson et al. 2008) and RLE, implemented in DESeq2 (Anders and Huber 2010; Love et al. 2014). However, neither of these methods employ any gene length normalization (their aim being to identify differentially-expressed genes between samples and thus they assume that the gene length is constant across samples. TPM (Transcripts Per kilobase Million) normalization (Li et al. 2008) extends the previously used RPKM (Reads Per Kilobase per Million reads) for single-end sequencing protocols (Mortazavi et al 2008) and its paired-end counterpart, FPKM (Fragments Per Kilobase per Million reads) (Trapnell et al. 2008), as both RPKM and FPKM proved to be inadequate and biased (Bullard et a1. 2010; Olshack et al, 2009; Wagner et al 2012). TPM employs a simple normalization scheme, where the raw read counts of each gene are divided by the gene length in kb and the total sum of all RPK is considered the library size of that sample. Thus TPM can be used for fully-elucidated genomes and for partial transcriptomes. Finally, the library size is divided by a million, and that number is employed as the scaling factor to scale each genes’ RPK value.

Under ideal conditions, a normalization methodology should account for all the major sources of error in a sample and should yield a dataset on which both between-sample and within-sample analyses can be performed. Smid et al. (2018) aimed to do this in their implementation of GeTMM (Gene length corrected trimmed mean of M-values) which combines gene-length correction with the normalization procedure TMM. GeTMM performs similarly TPM in intersample analyses but has clear advantages in intrasample comparisons (Smid et al. 2018).

Recent studies have shown that, in plants, gene expression is highly tissue specific and often varies with tissue age and the physiological status of the plant and the exact experimental conditions and, to date, there has not been any clear report of universal reference genes (Kozera and Rapacz 2013; Joseph et al 2018; Hong et al. 2008; Gutierrez et al. 2010). Coupled with recent findings that commonly-employed housekeeping genes may be far more variable in their expression than previously realized. Historically, selection of reference genes for qPCR studies has typically been arbitrary, with genes such as *25S* and *18S* rRNAs, *GAPDH*, and *Actin* commonly being selected without experimental validation (this being true for both plant and animal studies). In concert, these genes were often employed with the assumption that they are stably expressed across tissues. However, we now know that in many instances these commonly used RGs exhibit tissue and treatment specific variability (Chari et al., 2010; De Jonge et al., 2007). A previous preliminary study on a number of human cell lines and tumour versus matched normal tissue samples demonstrated that inappropriate choice of RGs may lead to errors when interpreting experiments involving quantification of gene expression (Janssens et al. 2004).

In an attempt to correct for over-expressed genes with variable expression edgeR employs the Trimmed Means of M-values (TMM) (Robinson and Olshack 2010) in which highly expressed genes and those that have a large variation of expression are excluded, whereupon a weighted average of the subset of genes is used to calculate a normalization factor. In the edgeR implementation precision (inverse of variance) weights are used to account for the fact that log fold changes from genes with higher read counts have lower variance on the logarithm scale. This typically excludes very highly expressed transcripts such as 25S and 18S ribosomal RNAs along with other very highly expressed transcripts from the initial transcript pool.

Many normalization methodologies assume that, depending on whether genes or transcripts are the base unit, that the length of this base unit is the same between samples. In diploids, this can be assumed to be correct at the gene level. However in organisms with high ploidy allelic variants of genes make this unlikely. If the experiment is being performed at the transcript level, due to alternate transcripts (particularly tissue specific alternate transcripts) the assumption does not hold at all. Thus corrections for transcript/gene lengths are required.

Panicum virgatum (switchgrass) is a tall, upright, bunchgrass that is a feature of North American prairies. *Panicum virgatum* is seen as potentially being an important bioenergy crop. It is an outcropping species that is an hybrid of two ancestral species (Lovell et al. 2021). Tetraploid P. virgatum plants have variously hybridized to generate disparate octoploid forms (Triplett et al 2012).

We extend the GeTMM implementation by removing over and under-expressed transcripts with edgeR and develop a novel PERL implementation to apply effective transcript length (including alternate transcript variants), GC skew and library lengths as additional normalization variables and apply this to public leaf RNA-seq datasets in *Panicum virgatum* cultivars. Being highly polyploid and with a newly available high quality genome sequence (Lovell et al 2021) and numerous high depth RNA-seq datasets available, many with multiple replicates Panicum virgatum makes an excellent reference test species for the software.

## Materials and Methods

### Identification of Switchgrass Datasets

Analyses on algorithm efficiency were performed using the tetraploid *Panicum virgatum* AP13 v5.1 (Lovell et al. 2021) genome as a reference. All transcripts (not just primary transcripts) were exported from Phytozome v 13 (Goodstein et al. 2012). Two datasets with three replicates apiece were employed for the analyses of differential expression. These were *Panicum virgatum* cv Alamo leaf day 9: SRA accessions SRR12851488; SRR12851477; SRR12851469 and *Panicum virgatum* cv Alamo leaf 5 months: SRA accessions SRR6485351; SRR6485352; SRR6485353 (Chen et al 2020).

### Read Pre-processing and Mapping

Prior to mapping, the reads to the reference transcripts polyA tails were manually clipped from transcripts (where they occurred).

The 9-day leaf was employed as control and the 5-month leaf samples were the test samples. For the SRA datasets, following Corchete et al. (2020) adapter removal only was performed with Trimmomatic 0.39 (LEADING:4 TRAILING:4 SLIDINGWINDOW:4:20MINLEN:50) (Bolger et al. 2014). However, instead of directly performing quality trimming with Trimmomatic reads were next error corrected with the error correction pipeline of SPAdes v 3.15.1 (Prjibelski et al. 2020). For paired end data, subsequent to SPAdes error correction the paired end data only was passed to Trimmomatic for quality trimming. For single end data all error corrected reads were employed as input for Trimmomatic.

Trimmed and error corrected reads were mapped to individual transcript sequences padded with Ns using HISAT2 v2.2.2.1 (Kim et al. 2019) mapped reads were enumerated with HTSeq v 0.12.4 (Anders et al. 2015) using a custom the Union approach. HISAT2’s native SAM output format was piped to SAMtools (Danecek et al. 2021) and output in BAM format. Mapped reads were analyzed for completeness of the transcript and the presence of missing exons, before being employed as input to our novel normalization algorithm.

### Data Export

Based on transcript sequences and read mappings, the following data were collected for input into the GeGC-TMM methodology: read lengths and insert sizes from the bam mapping file using samtools and picard tools; transcript sequence length and GC content using Emboss infoseq and geecee (Rice et al. 2000); count of mapped reads using Samtools.

### Algorithm Implementation

#### GeGC-TMM methodology for normalizing RNA-seq data

Transcripts were trimmed of adapter sequences and low-quality sequence regions using Trimmomatic (ref). Trimmed sequences were error corrected using the error correction portion of the SPAdes (ref) assembler pipeline.

GC content is another major factor that requires normalization. Risso et al. (2011) demonstrated that full quantile normalization is the most appropriate approach. For this methodology genes are stratified according to GC-content, with the normalized expression measures defined as:

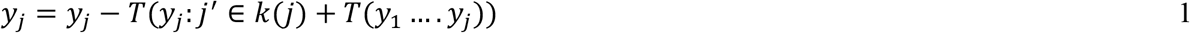

where *k*(*j*) denotes the GC-content stratum to which gene *j* belongs and *T* denotes the upper-quantile for control genes. The quantiles of the read count distributions are then matched between GC-bins, by sorting counts within bins and then taking the median of quantiles across bins.

Prior to GeTMM normalization, GC normalization was performed in the Bioconductor R package EDASeq package (Risso et al. 2011) using the withinLaneNormalization and betweenLaneNormalization methods for inter-sample and intra-sample normalization, respectively.

Outputs from EDASeq were input into the GeGC-TMM application, which is described below:

For normalization, the overall methodology is based on the work of Smid et al. (2018) for data normalization and the work of Corchete et al. (2020) for the stability ranking of genes. Below is a full mathematical treatment:

Define *Y*_*gk*_ as the observed count of mapped raw reads for gene *g* in library *k*. *μ*_*gk*_ is the true and *unknown* expression level (total number of transcripts) and *L*_*g*_ is the length of gene *g* and *N*_*k*_ is the total number of reads in library *k*.

For sequencing data, the gene-wise log-fold changes are defied as:

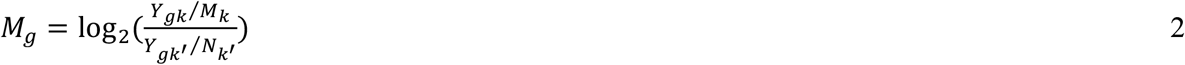

the corresponding absolute expression levels are:

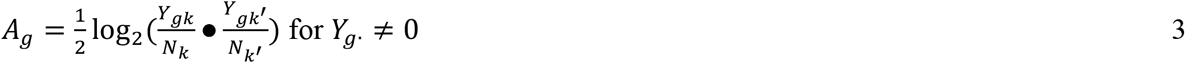

M and A values are employed to trim those genes/transcripts with too high (high expression and high variance) and too low (incomplete coverage). In our implementation we trimmed 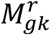 (sample k relative to sample r (where r is the reference set) for gene g) by 20% and the absolute expression, Ag, by 5%.

Thus, the final set of genes G* is a subset of the initial set G (ie *G* ∈ *G*). The above is implemented in the edgeR package (Robinson and Oshlack 2010) and this was employed for the initial stages of analysis.

The normalization factor for a sample k relative to a sample r is obtained as:

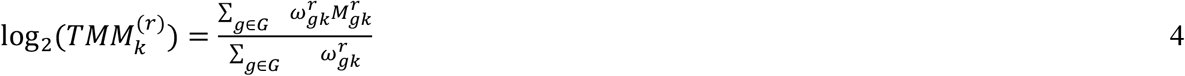

Where:

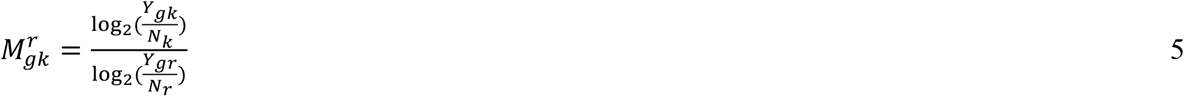

And:

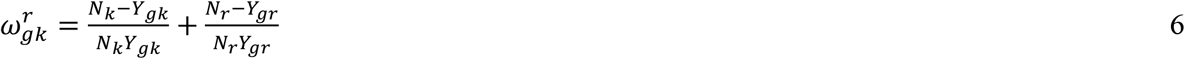

There is an additional implicit trimming here as *Y*_*gk*_ and *Y*_*gr*_ must be greater than 0.

For GeTMM normalization, mapped read data are first converted to RPK (reads per kilobase) and are scaled with the TMM scale factor, as defined above.

Converting raw read counts to RPK is as simple as dividing a gene (or transcript’s) raw read mapping count by the gene length in kilobases.

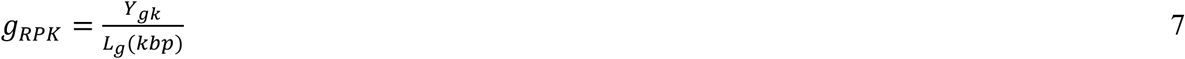

However, rather than the full length of the gene (or transcript of interest) it is better to use 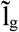, the Effective

Length of the feature of interest which is defined thus:

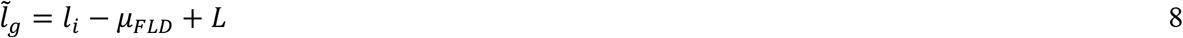

l_i_ is the length of the feature of interest, μ_FLD_ is the mean of the fragment length distribution, as determined from the mapped reads and ℒ is the sequence bias (if the mapping technique provides it). If ℒ is not provided, it is typically set to 1. For species without a reference genome and only an incomplete transcriptome

RPK scale factor is given as:

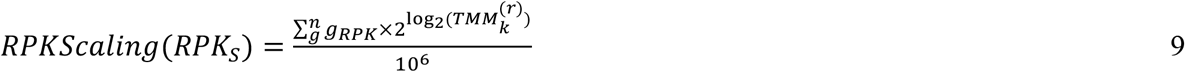

Where n is the total number of genes in G** (G** being the set of all genes after edgeR and GeTMM trimming).

The normalized read count for gene g thus becomes:

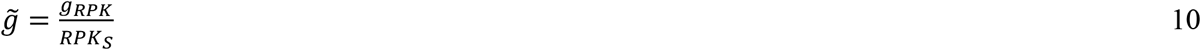

Where 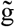 represents the GeTMM normalized gene count.

### Identification of Stably-expressed Genes

Corchete et al (2020) in their analysis of best in breed methodologies for RNA-seq based procedures for gene expression quantitative analyses advocated the use of coefficient of variance for ranking gene expression stability.

By their definition,

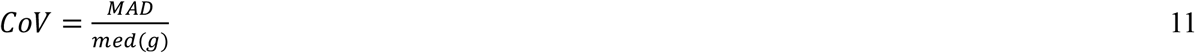

Where med(g) is the median value for gene ‘g’ and MAD is the median absolute deviation as given by:

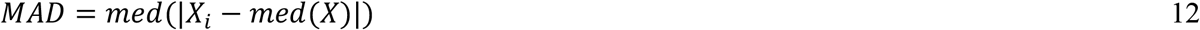

Where X is the xth gene and ‘i’ is the i’th sample and, in terms of algorithmic implementation:

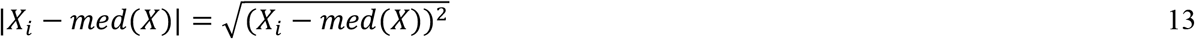

By calculating the CoV for each gene, the genes can be ranked in order of stability.

The full methodology and implementation methodology with how to run the code is given in the code itself (open source) and the accompanying documentation.

### Differential Expression Analysis

After running the RNA-seq mapped data through the GeGC-TMM application fold change was estimated from the edgeR (Robinson et al. 2010) regression model fit. To compare the cumulative effects of the different normalization protocols each method was applied in sequence and compared with the results of the full pipeline.

## Results

As the implementation of the GC-GeTMM is step-wise output from the individual steps can be exported for analysis. The RNA-seq data for nine day old and five month old plants were normalized independently. Each dataset had three replicates. Normally the normalized replicates would be averaged prior to determination of the coefficient of variation.

In this case, all datapoints were size sorted and the log_2_ of the length was determined. Fold change was determined with edgeR and log_2_ of fold change was plotted against log_2_ of transcript length (Figure 1). For each transcript length the mean fold change was plotted in red.

**Figure 1.**
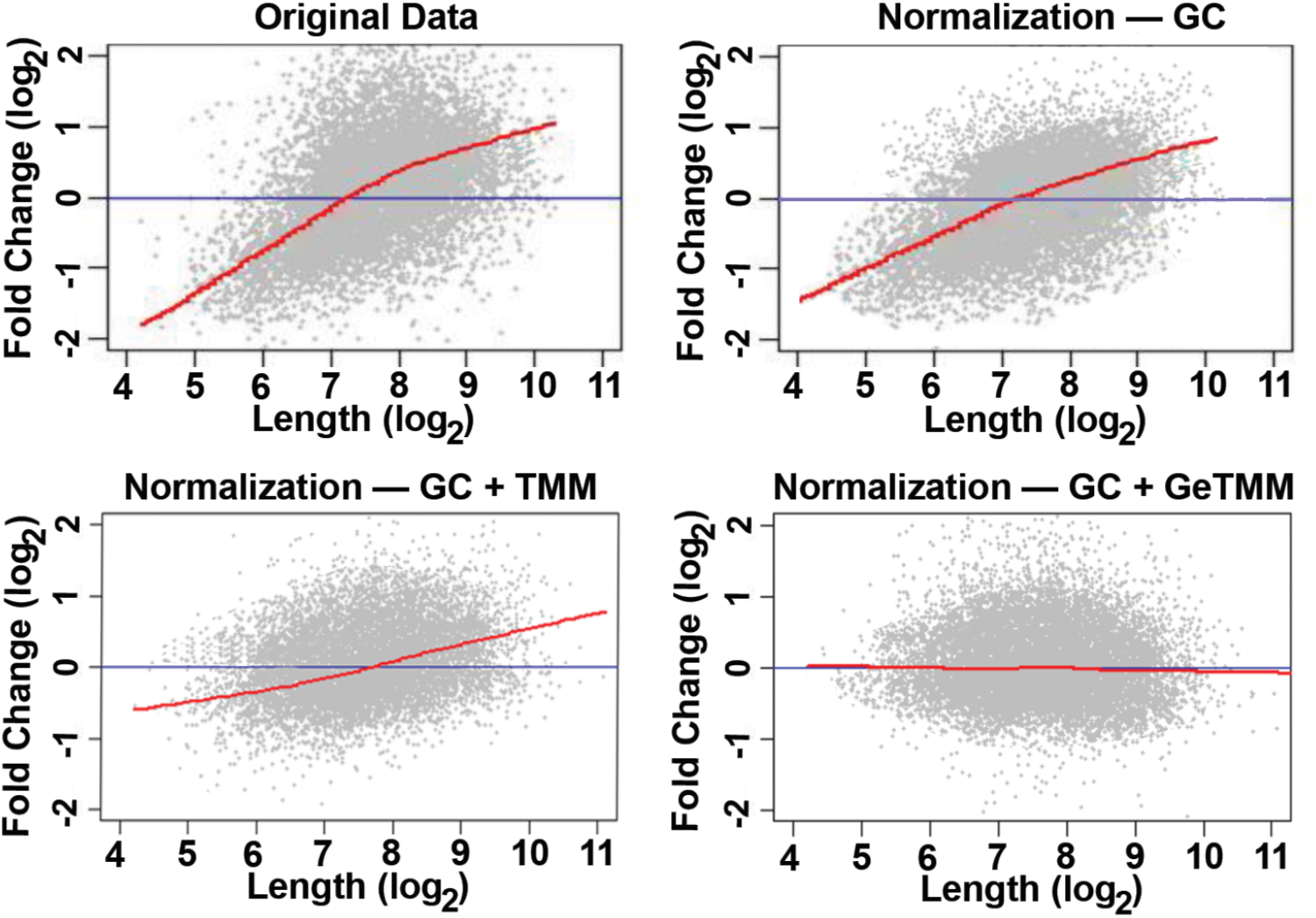
Analysis of the effects of different normalization methodologies on the quality of differential expression studies for *Panicum virgatum* 9 day old and 5 month old leaves. Top left: comparison of fold change against transcript length for the original data. Top right: effect of GC normalization on data quality. Bottom left: effect of TMM normalization on the GC normalized data. Bottom right: effect of combining GC normalization with GeTMM normalization.

Each step in the normalization process (GC, TMM, GeTMM) yields improvements in the data quality as the midline curve (red in Figure 1) tends towards the X-axis.

Subsequent to normalization, the replicates were merged and CoV values were determined (Table 1).

**Table 1.**
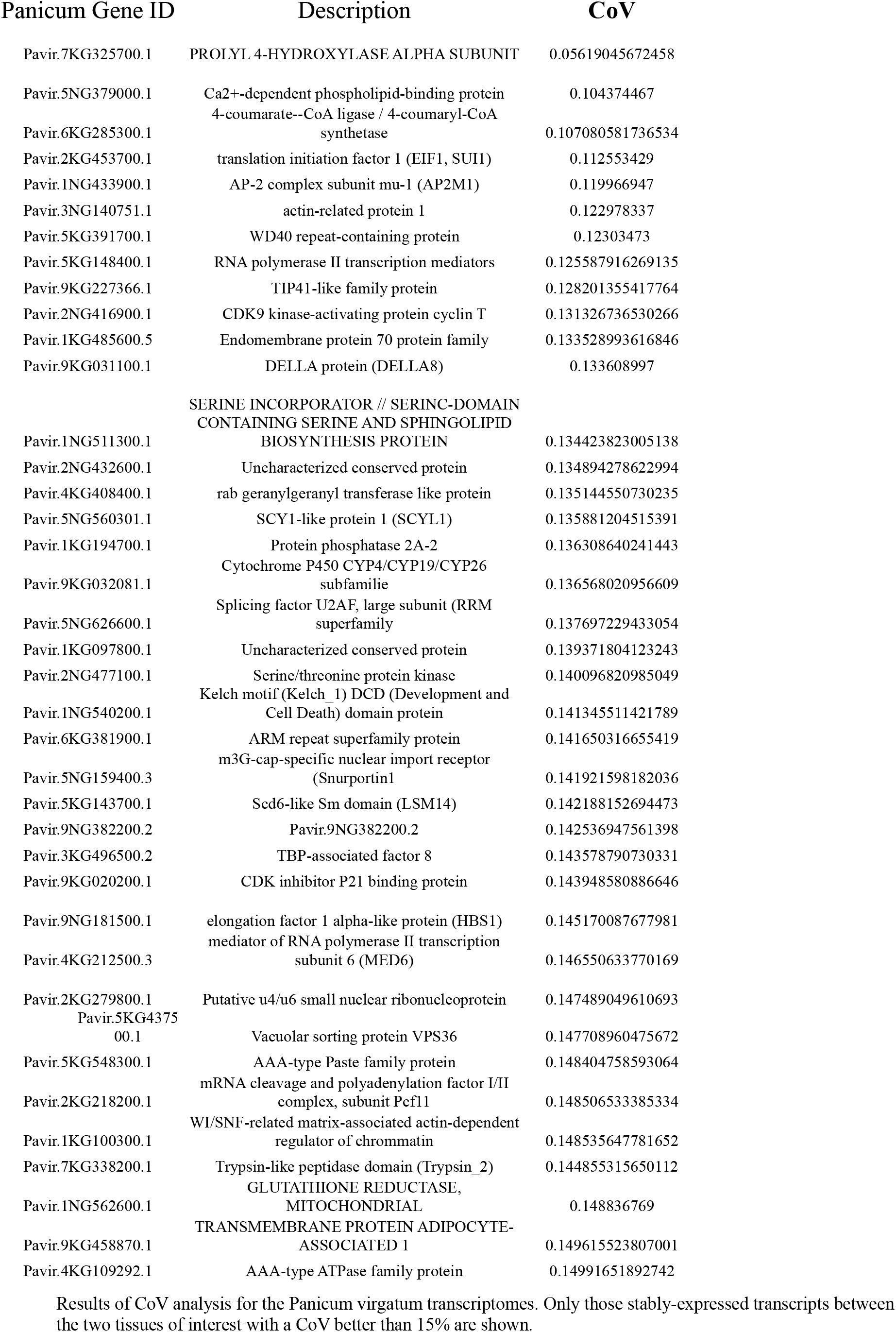
Results of CoV analysis for the Panicum virgatum transcriptomes. Only those stably-expressed transcripts between the two tissues of interest with a CoV better than 15% are shown.

CoVs were determined for all the transcripts at the genome (apart from those transcripts that failed TMM analyses as being too highly and unstably expressed). Focussing at the transcript level also allows differentially expressed and orthologous transcripts from the two founding genomes of *P. virgatum* to be analyzed. The results of the analysis are presented in Table 1.

## Discussion

The quest for stably expressed genes in multiple tissues and developmental stages is essential for the normalization of gene expression analyses by qRT-PCR and microarray studies. qRT-PCR is currently accepted as the gold standard for expression analyses, but is generally expensive and time-consuming. Microarray analyses are very high throughput but can suffer from issues of sensitivity. RNA-seq can afford a middle way. Many datasets are readily available from NCBI’s sequence read archive (SRA) and individual RNA-seq datasets can be generated for a few thousand dollars.

RNA-seq allows rapid mapping of reads to transcripts and the quick ranking of stable transcripts. Thus the GCGe-TMM application was developed to extend the Ge-TMM to correct GC bias along with transcript length bias. The application presented in this paper can simultaneously error correct RNA-seq data for import into other applications for differential expression analysis as well as ranking transcripts in terms of covariant of expression.

Figure 1 demonstrates that the three steps of normalization employed (GC bias, TMM and length bias) all significantly improve the error profile of the data, making it suitable for further analyses. For whole genome analyconses (Table 1) a conservative cutoff of 15% was chosen (typical cutoffs range between 30 and 40% (refs). As the methodology for TMM excludes highly-expressed transcripts but unstably expressed transcripts. This excludes some of the common highly-expressed transcripts (25S rRNA, 16S rRNA, GAPDH, TATA binding protein (Thellin et al 1999; Vandesompele et al 2002).

Improved normalization leads to improved and more reliable differential expression analyses.

## Conclusion

The GCGeTMM methodology presented in this paper affords improvements over the Ge-TMM implementation, particularly for monocot plants, with their generally higher GC content (Li and Du 2014).

The software described herein enables the simultaneous identification of stably expressed genes/transcripts and the identification of differentially-expressed transcripts at a whole genome and a gene/transcript level. It is applicable to species with complete genomes, but can also be used for those species without a genome (but with a transcriptome) as such it can be employed for orphan and under-funded species as it relies on transcript rather than gene level analyses. It can also be employed for subsets of transcripts (particularly for the analysis of stably expressed transcripts.

As such, the application presents a major step forwards for the analysis of stably expressed genes and differentially expressed genes in RNA-seq based gene expression analysis for plants, polyploids and those species with only partial or unsequenced genomes.

## Code and Data Availability

All data employed in this paper are publicly available. All code has been deposited in GitHub: https://github.com/gwydion1/expression.

